# *CXCR4* involvement in neurodegenerative diseases

**DOI:** 10.1101/181693

**Authors:** Luke W Bonham, Celeste M. Karch, Chun C. Fan, Chin Hong Tan, Ethan G. Geier, Yunpeng Wang, Natalie Wen, Iris J. Broce, Yi Li, Matthew J. Barkovich, Raffaele Ferrari, John Hardy (International FTD-GWAS Consortium (IFGC)), Parastoo Momeni, Günter Höeglinger, Ulrich Müller, Christopher P. Hess, Leo P. Sugrue, William P. Dillon, Gerard D. Schellenberg, Bruce L. Miller, Ole A. Andreassen, Anders M. Dale, A. James Barkovich, Jennifer S. Yokoyama, Rahul S. Desikan

## Abstract

Neurodegenerative diseases likely share common underlying pathobiology. Although prior work has identified susceptibility loci associated with various dementias, few, if any, studies have systematically evaluated shared genetic risk across several neurodegenerative diseases. Using genome-wide association data from large studies (total n = 82,337 cases and controls), we utilized a previously validated approach to identify genetic overlap and reveal common pathways between progressive supranuclear palsy (PSP), frontotemporal dementia (FTD), Parkinson’s disease (PD) and Alzheimer’s disease (AD). In addition to the *MAPT* H1 haplotype, we identified a variant near the chemokine receptor *CXCR4* that was jointly associated with increased risk for PSP and PD. Using bioinformatics tools, we found strong physical interactions between *CXCR4* and four microglia related genes, namely *CXCL12*, *TLR2, RALB* and *CCR5.* Evaluating gene expression from post-mortem brain tissue, we found that expression of *CXCR4* and microglial genes functionally related to *CXCR4* was dysregulated across a number of neurodegenerative diseases. Furthermore, in a mouse model of tauopathy, expression of *CXCR4* and functionally associated genes was significantly altered in regions of the mouse brain that accumulate neurofibrillary tangles most robustly. Beyond *MAPT*, we show dysregulation of *CXCR4* expression in PSP, PD, and FTD brains, and mouse models of tau pathology. Our multi-modal findings suggest that abnormal signaling across a ‘network’ of microglial genes may contribute to neurodegeneration and may have potential implications for clinical trials targeting immune dysfunction in patients with neurodegenerative diseases.

## INTRODUCTION

Uncovering the shared genetic architecture across neurodegenerative diseases may elucidate underlying common disease mechanisms and promote early disease detection and intervention strategies. Progressive supranuclear palsy (PSP), frontotemporal dementia (FTD), Parkinson’s disease (PD) and Alzheimer’s disease (AD) are age-associated neurodegenerative disorders placing a large emotional and financial impact on patients and society. Despite variable clinical presentation, PSP, AD and FTD are characterized by abnormal deposition of tau protein in neurons and/or glia in the frontal, temporal, and insular cortical gray matter and hindbrain^1,2^. PSP is associated with 4-repeat (4R) tau inclusions^1,3^; FTD is characterized by 3-repeat (3R)-only, 4R-only, or mixed (3R and 4R) tau inclusions or TAR DNA-binding protein 43 (TDP-43) aggregates^4^; and AD is characterized by extracellular amyloid plaques and neurofibrillary tangles (NFTs) composed of hyperphosphorylated tau (mixed 3R and 4R tau inclusions)^5,6^. While PD is classically characterized by alpha-synuclein deposits, recent studies support the role of tau and NFTs in modifying PD clinical sympotomatology and disease risk^7–9^. Prior work has shown that PSP shares overlapping pathobiology with FTD, AD, and PD^10–13^.

Genome-wide association studies (GWAS) and candidate gene studies have identified single nucleotide polymorphisms (SNP) within the H1 haplotype in *MAPT* locus (which encodes tau) that increase risk for PSP, FTD, AD, and PD^14–19^. However, beyond *MAPT*, the extent of genetic overlap across these diseases and how its relationship with common pathogenic processes observed in PSP, FTD, AD, and PD remains poorly understood.

Genomic studies evaluating shared risk among numerous phenotypes or diseases suggest genetic pleiotropy, where a single gene or genetic variant may impact different traits^17-20^. The recent proliferation of GWAS data for rare disorders like PSP paired with large studies of more common diseases such as AD, PD, and FTD provides unique opportunities to gain statistical power and identify risk loci which may not have otherwise been identified in the original GWAS study. Further, it allows for the systematic evaluation of genetic overlap across different disorders and thereby informing shared biological pathways and processes commonly altered in both conditions. Here, using previously validated methods^20–23^, we assessed shared genetic risk across PSP, PD, FTD, and AD. We then applied molecular and bioinformatic tools to elucidate the role of these shared risk genes in neurodegenerative diseases.

## METHODS

### Participant Samples

We obtained publicly available PSP-GWAS summary statistic data from the NIA Genetics of Alzheimer’s Disease Storage Site (NIAGADS), which consisted of 1,114 individuals with PSP (cases) and 3,247 controls (stage 1) at 531,451 SNPs (Table 1, for additional details see ^18^). In this study, we focused on stage 1 of the PSP GWAS dataset. Individuals were diagnosed with PSP according to NINDS criteria^24^. We evaluated complete summary statistic GWAS data from clinically diagnosed FTD, AD and PD. The International Parkinson’s Disease Genetics Consortium (IPDGC) provided PD-GWAS summary statistic data. The IPDGC cohort consists of 5,333 cases and 12,019 controls with genotype or imputed data at 7.689,524 SNPs (Table 1, for additional details see ^9^). The International FTD GWAS Consortium (IFGC) provided phase 1 FTD-GWAS summary statistic data, which consisted of 2,154 FTD cases and 4,308 controls with genotypes or imputed data at 6,026,384 SNPs (Table 1, for additional details see ^25^ and Supplemental Information). The FTD dataset included multiple subtypes within the FTD spectrum: bvFTD, semantic dementia, progressive non-fluent aphasia, and FTD overlapping with motor neuron disease. We obtained publicly available, AD-GWAS summary statistic data from the International Genomics of Alzheimer’s Disease Project (IGAP Stage 1) (see Supplemental Information). The IGAP Stage 1 cohort consists of 17,008 AD cases and 37,154 controls with genotyped or imputed data at 7,055,881 SNPs (Table 1, for additional details see ^26^ and Supplemental Information). All four cohorts were primarily of European ancestry and the studies’ authors controlled for population stratification using a principal components analysis approach. Inclusion/exclusion criteria were established by each study’s authors; please see the publications provided in Table 1 for additional details. In each study, informed consent was obtained from all subjects. Institutional approval was provided by each study’s respective committee.

**Table 1.** Pleiotropy Analysis Cohort Descriptions. Cohort descriptions with identifying details of the publishing study are provided.

| Disease/Trait | Total N | # SNPs | Reference |
| --- | --- | --- | --- |
| Progressive<br>Supranuclear Palsy<br>(PSP) – Phase 1 | 4,361 | 531,451 | Höglinger, G et al. Identification of common variants influencing risk of the tauopathy progressive supranuclear palsy. Nat Genet. 2011; 43; 699-705. |
| Frontotemporal<br>Dementia (FTD) –<br>IFGC phase 1 | 6,462 | 6,026,384 | Ferrari, R et al. Frontotemporal dementia and its subtypes: a genome-wide association study. Lancet Neuro. 2014;13;686-99. |
| Alzheimer’s disease<br>(AD) – Phase 1 | 54,162 | 7,055,881 | Lambert JC, Ibrahim-Verbaas CA, Harold D, et al. Meta-analysis of 74,046 individuals identifies 11 new susceptibility loci for Alzheimer's disease. Nat Genet. 2013;45:1452-8. |
| Parkinson’s disease<br>(PD) | 17,352 | 7,689,524 | Nalls, M.A. et al. Imputation of sequence variants for identification of genetic risks for Parkinson’s disease: a meta-analysis of genome-wide association studies. Lancet. 2011;377;641-9. |

### Identification of shared risk loci – conjunction FDR

We evaluated SNPs associating with PSP (Phase 1), FTD (IFGC Phase 1), AD (IGAP Stage 1) and PD (IPDGC Phase 1) using techniques for evaluating genetic pleiotropy (for additional details see Supplemental Information and ^21,23^). Briefly, for two phenotypes A and B, pleiotropic ‘enrichment’ of phenotype A with phenotype B exists if the effect sizes of the genetic associations with phenotype A become larger as a function of increased association with phenotype B. For each phenotype, Z scores were derived for each SNP association given the Wald statistics. We then corrected the Z scores for potential genomic inflation(for additional methodological details used in our study, see Supplemental Information and see ^27^ for additional information on genomic inflation). To assess enrichment, we constructed fold-enrichment plots of nominal –log_10_(p) values for all PSP SNPs and a subset of SNPs determined by the significance of their association with AD, FTD, and PD. Under expected null, the average effect sizes given a group of SNP would be the same when the conditioned effect increases. Therefore, in fold-enrichment plots, enrichment is indicated by an upward deflection of the curve for phenotype A if it shares genetic effects with phenotype B. To assess for polygenic effects below the standard GWAS significance threshold, we focused the fold-enrichment plots on SNPs with nominal –log_10_(p) < 7.3 (corresponding to p-value > 5x10^−8^). The enrichment can be interpreted in terms of true discovery rate (TDR = 1 – False Discovery Rate [FDR]) (for additional details see Supplemental Information and ^28^).

To identify specific loci involved in both PSP and AD, FTD or PD, we computed conjunction FDR^21–23^. Conjunction FDR, denoted by FDR_trait1& trait2_ is defined as the posterior probability that a SNP is null for either phenotype or both simultaneously, given the p-values for both traits are as small, or smaller, than the observed p-values. A conservative estimate of the conjunction FDR is given by the maximum statistic in taking the maximum of FDR_trait1|trait2_ and FDR _trait2|trait1_ (for additional details see Supplemental Information and ^21^). We used an overall FDR threshold of < 0.05. To visualize the results of our conjunction FDR analysis, we constructed Manhattan plots based on the ranking of conjunction FDR to illustrate the genomic location of the pleiotropic loci. Rather than representing novel risk variants where replication is needed in independent datasets, conjunction FDR pinpoints genetic variants *jointly* associated with two or more phenotypes/diseases and in this context, ‘replication’ may not be meaningful^21^.

### Functional evaluation of shared risk loci

To assess whether the PSP, FTD, AD, and PD overlapping SNPs modify gene expression, we evaluated *cis*-expression quantitative trait loci (eQTLs, DNA sequence variants that influence the expression level of one or more genes) in a publicly available dataset from 134 neuropathologically confirmed normal control brains (UKBEC, http://braineac.org/)^29^ and validated these eQTLs in the GTex dataset^30^. We also evaluated eQTLs using a blood-based dataset^31^. We applied an analysis of covariance (ANCOVA) to test for association between genotypes and gene expression. SNPs were tested using an additive model.

### Network based functional association analyses

To evaluate potential protein and genetic interactions, co-expression, co-localization and protein domain similarity for the pleiotropic genes, we used GeneMANIA (www.genemania.org), an online web-portal for bioinformatic assessment of gene networks^32^. In addition to visualizing the composite gene network, we also assessed the weights of individual components within the network^33^.

### Gene expression alterations in PSP, PD, and FTD brains

To determine whether pleiotropic genes were differentially expressed in PSP, PD, and FTD brain tissue, we analyzed gene expression of pleiotropic genes in publically available datasets. We analyzed gene expression data from: 1) the temporal cortex and cerebellum of 80 control and 84 PSP brains (syn5550404); 2) the frontal cortex; hippocampus and cerebellum of 11 controls and 17 FTLD-U (7 brains with or 10 brains without *progranulin* (*GRN*) mutations) (Gene Expression Omnibus (GEO) dataset GSE13162)^34^; and 3) the substantia nigra of 23 control and 22 PD brains (GEO dataset GSE7621)^35^.

### Evaluating gene expression of pleiotropic loci in tau transgenic mouse models

We evaluated gene expression profiles for the nearest genes associated with our shared risk loci using publicly available P301L-tau transgenic mouse model (mutant human *MAPT* gene) from mouseac (www.mouseac.org)^36^. Briefly, microarray gene expression data was collected from three brain regions (cortex, hippocampus and cerebellum) from wild-type and P301L-tau transgenic mice. Gene expression levels were log transformed and expressed as a function of age. The presence of NFT pathology was evaluated and scored as previously reported by immunohistochemistry^36^. Using repeated measures ANOVAs within the hippocampus, cortex and cerebellum, we examined whether gene expression levels of PSP, FTD, AD and PD variants are significantly different between the P301L-tau transgenic and wild-type mice, across 2, 4, 8 and 18 months of age. To maximize our ability to detect an effect, we used all expression data available for each line of mice and age grouping. As our data was publicly available, there was no randomization or blinding of the data. Please see (www.mouseac.org)^36^ for exact sample sizes used in each analysis and additional information on the mouse data used in this study.

### Code Availability

The code used to conduct pleiotropy analyses is not yet publicly available. Please contact the authors with any inquiries related to the code.

## RESULTS

### Selective shared genetic risk between PSP, PD, and FTD

We observed SNP enrichment for PSP SNPs across different levels of significance of association with FTD (Figure 1a). Using progressively stringent p-value thresholds for PSP SNPs (i.e. increasing values of nominal –log_10_(p)), we found up to 150-fold genetic enrichment using FTD and lower, but still notable, enrichment in PD (Figure 1a-b). In contrast, we found minimal or no enrichment in PSP SNPs as a function of AD (Figure 1a-b).

**Figure 1.**
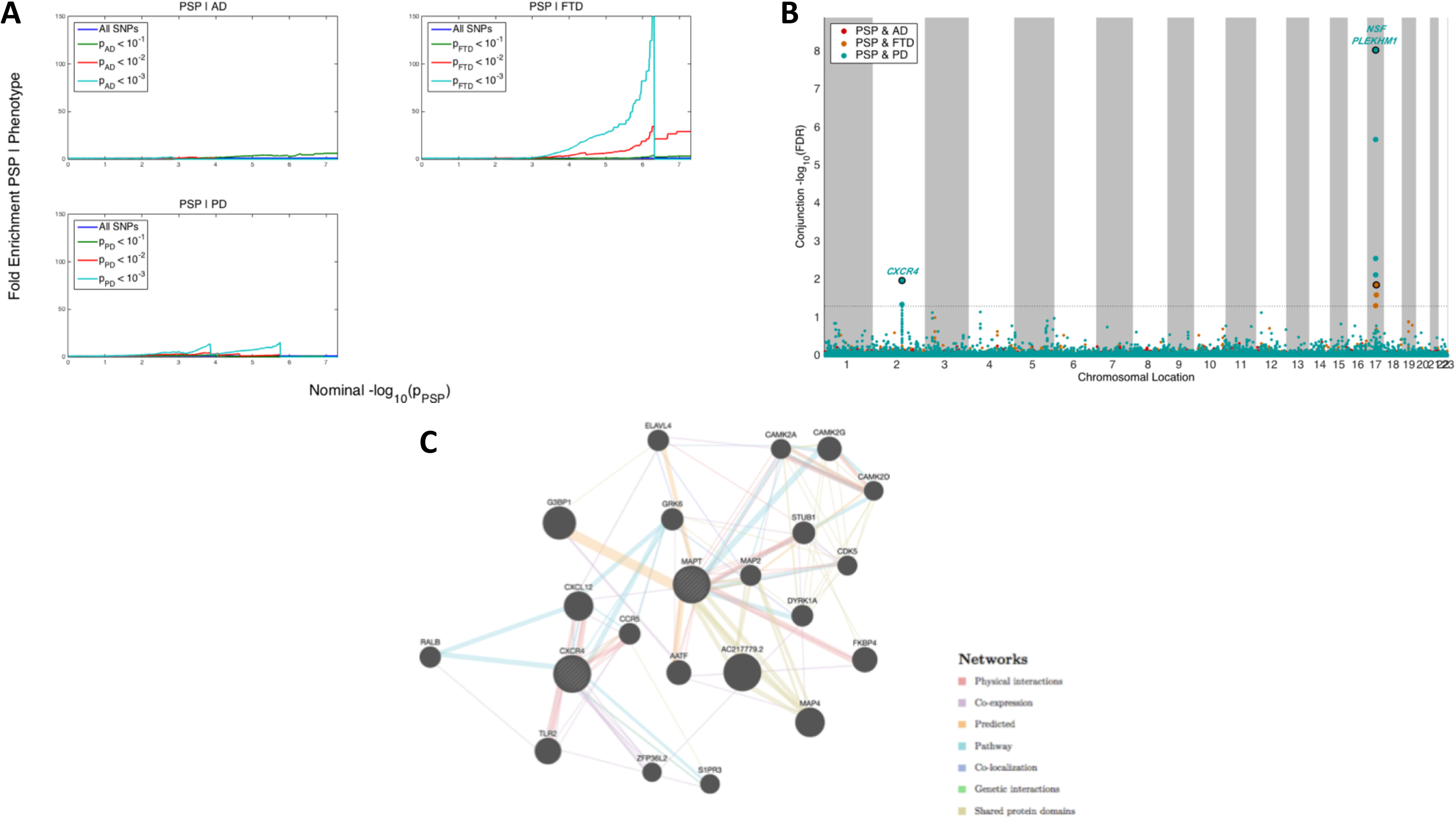
Shared genetic risk across PSP, AD, FTD and PD. **(a)** Fold enrichment plots of enrichment versus nominal −log10 p-values (corrected for inflation) in progressive supranuclear palsy (PSP) below the standard GWAS threshold of p < 5x10^−8^ as a function of significance of association with Alzheimer’s disease (AD, panel A), frontotemporal dementia (FTD, panel B) and Parkinson’s disease (PD, panel C) at the level of −log10(p) ≥ 0, −log10(p) ≥ 1, −log10(p) ≥ 2 corresponding to p ≤ 1, p ≤ 0.1, p ≤ 0.01, respectively. Blue line indicates all SNPs. **(b)** ‘Conjunction’ Manhattan plot of conjunction and conditional –log10 (FDR) values for progressive supranuclear palsy (PSP) (black) given Alzheimer’s disease (AD; PSP|AD, red), frontotemporal dementia (PSP; PSP|FTD, orange) and Parkinson’s disease (PD; PSP|PD). SNPs with conditional and conjunction –log10 FDR > 1.3 (i.e. FDR < 0.05) are shown with large points. A black line around the large points indicates the most significant SNP in each LD block and this SNP was annotated with the closest gene, which is listed above the symbols in each locus. **(c)** Network interaction graph illustrating physical interactions, co-expression, predicted, pathway, co-localization, gene interactions and shared protein domains for *CXCR4* and *MAPT*. Network plot was generated from GeneMANIA (www.genemania.org).

At a conjunction FDR < 0.05, we identified 2 SNPs that were associated with increased risk for both PSP and PD (Table 2, Figure 1b): rs749873 (intergenic; closest gene = *CXCR4*, on chr2, minimum conjunction FDR = 0.01; Supplemental Figure 1a) and rs11012 (UTR-3; closest gene = *PLEKHM1* within *MAPT* region on chr 17, PSP p-value = 4.9 x 10^−39^, minimum conjunction FDR =9.3 x 10^−9^). *CXCR4* is a chemokine receptor implicated in immune processes, microglia recruitment, neuronal guidance, neural stem cell proliferation, and neurodevelopmental processes^37–40^. At a conjunction FDR < 0.05, we identified 1 SNP that was associated with increased risk for both PSP and FTD (Table 2, Figure 1b): rs199533 (exonic; closest gene = *NSF* within *MAPT* region on chr 17, PSP p-value = 3.5 x 10^−41^, minimum conjunction FDR =9.3 x 10^−9^; Supplemental Figure 1b). Notably, rs11012 and rs199533 is in strong linkage disequilibrium (LD) with SNP rs1800547, which tags the H1 haplotype of *MAPT* (rs11012 pairwise D’=0.96, r^2^=0.71; rs199533 pairwise D’=1, r^2^=0.94). The H1 haplotype of *MAPT* has been implicated in risk for both PSP, PD, and FTD^18,19^. In contrast to PSP and PD as well as PSP and FTD, we found no significant overlapping loci between PSP and AD.

**Table 2.** Pleiotropy Analysis Results. Results from pleiotropy analyses are shown. For each SNP, the nearest gene, chromosome, reference allele, alternative allele, conjunction p-value, conjunction analysis, and raw p-value from each of the 4 available GWAS is provided.

| rsID | Gene | Chromosome | Ref Allele | Alt Allele | Conjunction P-value | Conjunction Trait | PSP Raw P-value | PD Raw P-value | FTD Raw P-value | AD Raw P-value |
| --- | --- | --- | --- | --- | --- | --- | --- | --- | --- | --- |
| rs749873 | *CXCR4* | 2 | C | T | 0.01 | PD | 1.32E-04 | 6.66E-05 | NA | 0.48 |
| rs199533 | *NSF* | 17 | A | G | 9.27E-09 | FTD | 3.52E-41 | 1.07E-15 | 4.89E-05 | 0.01 |

### Expression quantitative trait loci (eQTL) and gene expression analyses

Consistent with our previous findings, rs749873 was associated with expression of *MCM6* within the *CXCR4* locus in human brains and with *CXCR4* expression in whole blood^41^. In the human CNS, rs749873 modifies MCM6 expression in the tibial nerve^41^. Interestingly, rs749873 is in high LD with rs2011946 (r^2^=0.91; D’=1), a SNP we previously reported to be shared across PSP and corticobasal degeneration (CBD)^41^. As previously reported, rs199533 was significantly associated with *MAPT* expression in human brains (p=2 x 10^−12^)^41^.

### Protein-protein and co-expression networks for MAPT and CXCR4

Using GeneMANIA, we examined the proteins that physically interact with and/or are co-expressed with *MAPT* or *CXCR4* (Figure 1c). As previously reported^41^, *CXCR4* demonstrated the strongest physical interaction with chemokine motif ligand 12 (*CXCL12)*, toll-like receptor 2 (*TLR2),* Ras-related protein-B *(RALB)* and C-C chemokine receptor 5 (*CCR5)* (Figure 1c) (Supplemental Table 1). *CXCL12* is the ligand for *CXCR4* ^42,43^. *TLR2* encodes a toll-like receptor which is utilized by the innate immune system to detect pathogenic material – it is expressed on microglia as well as astrocytes and has been proposed as an inhibitor of neural progenitor cell proliferation^44^. *RALB* is a small GTPase protein that interacts with *CXCL12* and is implicated in B-cell migration^45^. *CCR5* is a chemokine structurally related to *CXCR4* and modulates the migration of microglia and blood brain barrier integrity^46,47^.

We found that *MAPT* showed robust physical interactions with *FKBP4* and *STUB1* (Figure 1c; Supplemental Table 1). *FKBP4* (also known as FKBP52) is a peptidylprolyl isomerase which is involved in dynein interaction and glucocorticoid receptor movement to the nucleus^48^. *STUB1* is a ubiquitin ligase with diverse functions that has been implicated in AD^49^.

### CXCR4 gene expression alterations in PSP, PD and other neurodegenerative disease brains

We next sought to determine whether *CXCR4* is differentially expressed in PSP, PD, and FTD brains. We included FTD brain tissues in our analyses because no SNP data for rs749873 was available for primary pleiotropy analysis in FTD, leaving open the possibility that variation near *CXCR4* expression may be altered in FTD. Compared with control brains, we found that *CXCR4* was significantly upregulated in brains with a neuropathological diagnosis of PSP and FTD, especially within the cerebellum and hippocampus (Figure 2a-b, Table 3). Further, we found *CXCR4* was also significantly upregulated in PD cases (Figure 2c; Table 3). We also evaluated expression levels in PSP, FTD, and PD brains of the four genes that showed strong physical interactions with *CXCR4* in our network analyses (*CXCL12*, *TLR2, RALB* and *CCR5;* weighted connection with *CXCR4* > 0.25, see above). We found that *CXCL12* expression was significantly dysregulated in PSP, and PD, but not in FTD (Table 3). *TLR2* levels were significantly altered specifically in FTD. Of note, *CCR5* levels were not available for analysis. *MAPT* expression was not significantly altered in PSP, FTD or PD brains relative to controls (Table 3). Thus, our data suggest that expression of *CXCR4* and functionally associated genes are significantly altered in regions of the brain susceptible to different forms of neurodegenerative disease pathology.

**Table 3.** Gene expression analysis in PSP, PD, and FTD brains. P-values from the gene expression analyses in pathologically confirmed cases are shown. When multiple probes were available for each gene, the p-value for the first probe sorted by numerical order is provided. When multiple regions were used in an analysis, brain region was included as a covariate. For additional details on the probes and effect sizes, please see the Supplemental Material. FTD+GRN – frontotemporal dementia caused by *granulin* mutations. FTD-GRN – sporadic frontotemporal dementia. CRBL – Cerebellum; TC – Temporal cortex; SN-Substantia nigra; FC – frontal cortex; HIP – hippocampus.

|  | Diagnosis<br>(Cohort) | PSP |  | FTD+*GRN* | FTD-*GRN* | PD |
| --- | --- | --- | --- | --- | --- | --- |
|  | Tissue Type | CRBL | TC | FC, HIP,<br>CRBL | FC, HIP,<br>CRBL | SN |
|  | N | 84 | 84 | 7 | 10 | 22 |
| Gene<br>Name | CXCR4 | 6.31x10<sup>-4</sup> | 4.84x10<sup>-4</sup> | 1.86x10<sup>-4</sup> | 0.03 | 0.03 |
|  | CXCL12 | 0.03 | 5.22x10<sup>-4</sup> | 0.86 | 0.98 | 0.03 |
|  | RALB | 0.17 | 0.74 | 0.78 | 0.34 | 0.38 |
|  | TLR2 | 0.37 | 0.23 | 1.07x10<sup>-3</sup> | 0.02 | 0.85 |
|  | MAPT | 0.30 | 0.22 | 0.09 | 0.38 | 0.45 |
|  | TMEM119 | 0.74 | 0.39 | N/A | N/A | 0.27 |
|  | AIF1 | 0.08 | 0.68 | 0.12 | 0.82 | 0.48 |

**Figure 2.**
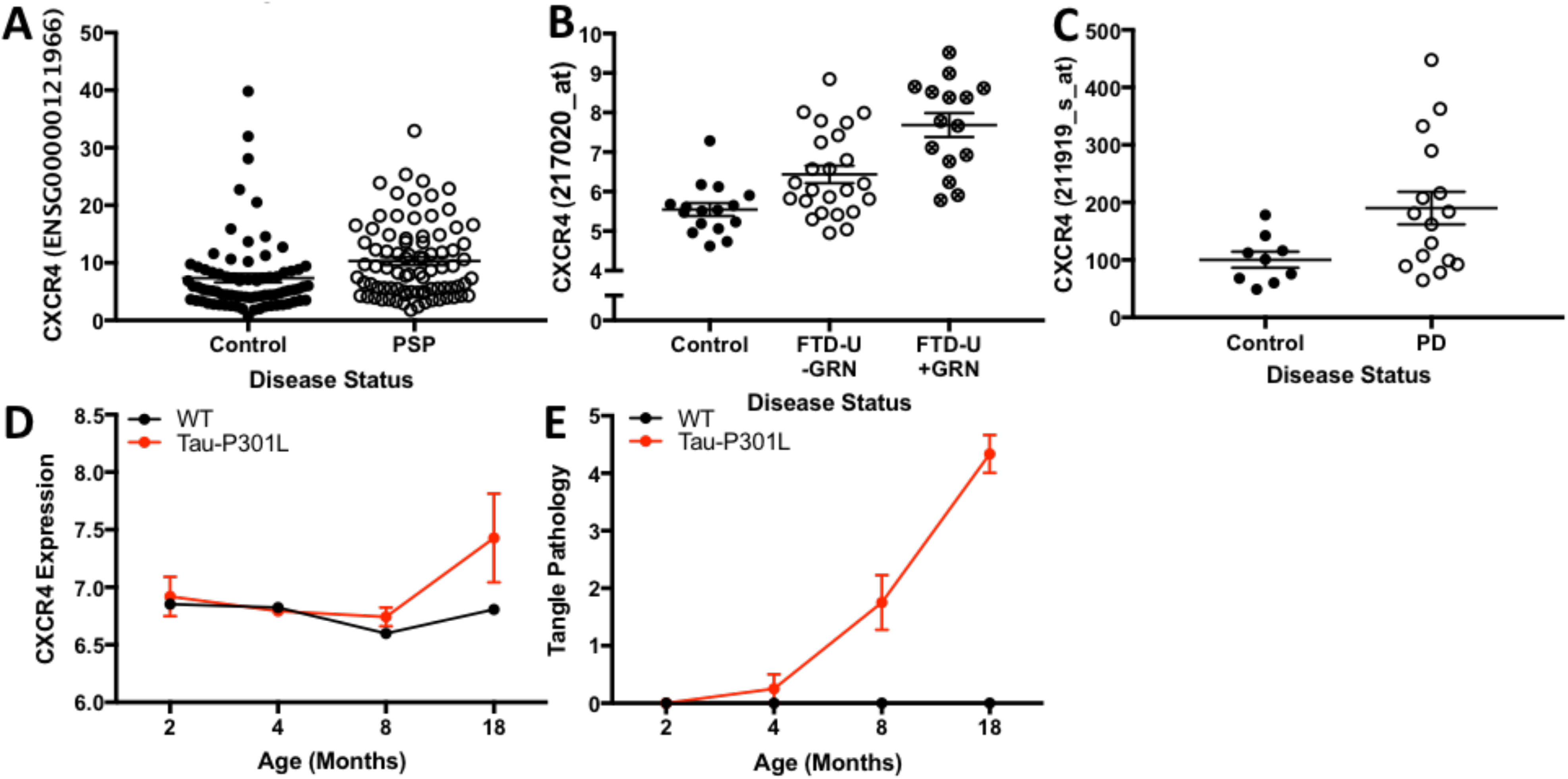
*CXCR4* is differentially expressed in human neurodegenerative diseases and in mouse models of tauopathies. **(a)** *CXCR4* expression in neuropathologically normal tissue compared to PSP. **(b)** *CXCR4* expression in neuropathologically normal tissue compared to sporadic FTD and FTD due to *GRN* mutations. **(c)** *CXCR4* expression in neuropathologically normal tissue compared to PD. **(d-e)** Line plots illustrating *CXCR4* gene expression in tau transgenic (red line) and wild-type mice (black line) from 2 to 18 months of age in the **(d)** hippocampus. **(e)** Total tau pathology over time is also illustrated.

Given the role of *CXCR4* in astroglial signaling and microglial activation^50^, we asked whether the observed upregulation of *CXCR4* in neurodegenerative disease was driven by proliferation and activation of microglia in response to neurodegenerative pathology. To assess this, we evaluated whether commonly used markers of microglial activation, namely *TMEM119* and *AIF1*^51,52^, are also upregulated across neurodegenerative diseases. Unlike *CXCR4*, across all evaluated brain regions within PSP, FTD, and PD brains, we found no evidence for altered *TMEM119* and *AIF1* expression (Table 3).

We next examined whether *CXCR4* and *AIF1* or *TMEM119* were significantly correlated in PSP, FTD, or PD brains. We observed a significant correlation between *CXCR4* and *AIF1* in FTD brains (combined FTD-GRN+ and FTD-GRN-: r^2^=0.35 and p<1x10^−4^). As indicated in Table 3, *TMEM119* was not available within the FTD dataset. In PSP brains, we found a marginally significant correlation between *CXCR4* and *TMEM119* (PSP *AIF1*: r^2^=5.7x10^−4^ and p=0.83; PSP *TMEM119*: r^2^=0.05 and p=0.05). However, *CXCR4* and *AIF1* or *TMEM119* were not correlated in PD brains (*AIF*: r^2^=9.15x10^−5^ and p=0.96 and *TMEM119*: r^2^=0.10 and p=0.10). Thus, these findings provide evidence suggesting that *CXCR4* could contribute to neurodegeneration through microglial activation as well as pathways beyond microglial dysfunction.

### CXCR4 expression is elevated in tau transgenic mouse model

To assess whether *CXCR4* expression is associated with NFT pathology, we examined *CXCR4* expression in brains from a transgenic mouse model of tau aggregation (Tau-P301L transgenic mice). Tau-P301L mice develop NFT pathology in the hippocampus and neocortical regions by 6 months of age, while the cerebellum fails to develop NFT pathology^36,53^. *CXCR4* expression was significantly elevated in the hippocampus (F = 10.3, p = 0.0023) in Tau-P301L mice compared with wild-type mice (Figure 2d). In contrast, within the cerebellum (F = 0.001, p = 0.98) and cortex (F = 1.06, p = 0.308), we found no difference in *CXCR4* expression between the Tau-P301L and wild-type mice across any of the time points. *MAPT* expression, however, was not significantly altered in Tau-P301L mice compared with wild-type in both the hippocampus and cerebellum (Supplemental Figure 2).

We additionally assessed expression levels of the four genes (*CXCL12*, *TLR2, RALB* and *CCR5*) that showed strong physical interactions with *CXCR4* in our network analyses (see Figure 1c). Within the hippocampus and cortex, *TLR2* was significantly elevated over time (F = 7.07, p = 1.35 x 10^−7^ and F = 2.9, p = 8.19 x 10^−6^, respectively) and *CCR5* expression was significantly decreased over time (F = 0.3, p = 0.002) in Tau-P301L mice but not in wild-type mice. Expression of *CXCL12* (F = 0.19, p = 0.06) and *RALB* (F = 0.02, p = 0.09) were not significantly altered in Tau-P301L mice (Supplemental Figures 3-6). Within the cerebellum, which remains free of tau aggregates, we found no evidence for gene expression alterations in *CXCL12*, *TLR2, RALB* and *CCR5* in Tau-P301L mice compared with wild-type mice. Thus, expression of *CXCR4* and functionally associated genes is significantly altered in regions of the mouse brain that accumulate NFTs most robustly.

Finally, we evaluated whether *TMEM119* and *AIF1* expression is associated with *CXCR4* expression and significantly elevated within the hippocampus. Across all evaluated time points (2, 4, 6 and 18 months) and within wild-type and Tau-P301L transgenic mice, we found that *TMEM119* and *AIF1* levels were significantly correlated with *CXCR4* expression specifically within the hippocampus (Supplemental Results). Similarly, predominantly within the hippocampus, we found that *TMEM119* and *AIF1* expression was elevated over time in Tau-P301L mice compared with wild-type mice (Supplemental Results). Evaluating *CXCR4* associated genes, we found a strong association between microglia and inflammation related *TLR2* and *CCR5* expression and *TMEM119* and *AIF1* expression, within the hippocampus, cortex and cerebellum (Supplemental Results). In contrast, we found no correlation between levels of *CXCL12* and *RALB* with *TMEM119* or *AIF1* (Supplemental Results). Together, these findings suggest that innate immune system signaling and microglial activation markers are associated with neurodegeneration and tauopathy.

## DISCUSSION

We identified *CXCR4* as a novel locus associated with increased risk for both PSP and PD. Building on extensive prior work, we also confirmed the role of variants within *MAPT* in driving PSP, PD, and FTD risk. We found that *CXCR4* and functionally associated genes exhibit altered expression across a number of neurodegenerative diseases. In a mouse model of tauopathy, *CXCR4* and functionally associated genes were altered in the presence of tau pathology. Together, our findings suggest that alterations in expression of *CXCR4* and associated microglial genes may contribute to age-associated neurodegeneration.

Utilizing GWAS summary statistics from multiple neurodegenerative diseases, our results suggest that shared genetic risk factors may underlie the pathobiological processes occurring in PSP and PD. We found up to 150-fold enrichment in PSP as a function of FTD, lower enrichment in PD, and no enrichment in AD. These findings were unexpected given the established role of the *MAPT* H1 haplotype in AD^26^. Despite the lack of strong genetic association across these three neurodegenerative diseases, we found that *CXCR4* expression was altered in brains that are pathologically confirmed for PSP, PD, and FTD. Thus, these findings support our hypothesis that these three neurodegenerative disorders share common pathobiological pathways.

*CXCR4* is a chemokine receptor protein with broad regulatory functions in the immune system and neurodevelopment^39,40,43,54–56^. *CXCR4* has been shown to regulate neuronal guidance and apoptosis through astroglial signaling and microglial activation^50^. Furthermore, it has been shown that *CXCR4* is involved in cell cycle regulation through p53 and Rb^57,58^. Importantly, small molecular agonists and antagonists to CXCR4 have been described^59^. AMD3100 is an FDA-approved CXCR4 antagonist and a CXCR7 agonist that is commonly used to enhance hematopoietic stem cell proliferation^60^.

Using a bioinformatics approach, we identified four genes, namely *CXCL12*, *TLR2, RALB* and *CCR5* that showed a strong association with *CXCR4*. We additionally found that expression of *CXCR4* and functionally associated genes were altered in multiple neurodegenerative diseases and associated with hippocampal tau pathology in transgenic mouse models. These findings suggest that a network of *CXCR4* and associated genes may act in concert to influence neurodegeneration.

Given that PSP, PD, and FTD are neuropathologically characterized by degeneration in the midbrain, cerebellum, and (to a lesser extent) neocortical regions^1,2,61^, our findings may suggest that subtle alterations of *CXCR4* and *MAPT* may predispose to regionally specific brain degeneration in later life. Though we found no evidence of altered expression of known microglial markers within human neurodegenerative brains, we found that *CXCR4* expression was significantly upregulated in PSP, FTD, and PD brains. Furthermore, within our transgenic tauopathy mouse model, we found a strong association between *CXCR4* and *TMEM119* and *AIF1* within the hippocampus. Additionally, across all evaluated regions, we found a strong relationship between microglia and inflammation related *CCR5* and *TLR2* expression and *TMEM119* and *AIF1* expression. Finally, expression of *TMEM119* and *AIF1* was markedly elevated predominantly within the hippocampus in Tau-P301L mouse model expression data. Thus, results from the Tau-P301L mouse model data suggest that upregulated *CXCR4* expression observed during age-associated neurodegeneration may be related to inflammatory mechanisms.

Our study benefits from its use of multiple well-validated GWAS datasets and its integration of multiple information modalities ranging from population level genetic data to RNA expression in mouse models of tauopathy. Our study used GWAS data composed of common SNPs and thereby cannot inform the potential role that rare variation may play as a risk factor for PSP, PD, and FTD. A limitation of our study is that we do not have records indicating which GWAS SNPs were directly genotyped versus imputed, limiting our ability to assess the quality of SNP ascertainment in our cohorts. Further, while we demonstrated that rs749873 modified *CXCR4* expression in human blood, we were unable to test for an eQTL in large dataset of control human brains. Given differences in sample size between the two tissue types, this could be a function of statistical power. Indeed, previously work has shown that many eQTLs are shared across tissues and that ability to detect an eQTL varies with sample size^30^.

In conclusion, by integrating large neurodegenerative GWAS data with gene expression data from neurodegenerative diseases and transgenic mouse models, our multi-modal findings indicate that *CXCR4* is associated with PSP and PD neurodegeneration. Clinically, our results provide additional evidence that immune and microglial dysfunction contribute to the pathophysiology in PSP, PD, and FTD. These findings have important implications for future work focused on monitoring microglial activation as a marker of disease progression and on developing anti-inflammatory therapies to modify disease outcomes in patients with neurodegenerative diseases.

## ACKNOWLEDGEMENTS

We thank the International FTD-GWAS Consortium (IFGC), International Parkinson’s Disease Genomic Consortium (IPDGC) and International Genomics of Alzheimer’s Project (IGAP) for providing summary statistics data for these analyses. Further acknowledgments for IFGC, IPDGC and IGAP are found in the Supplemental material. This research was supported by grants from the National Institutes of Health (NIH-AG046374 [CMK] and K01 AG049152 [JSY]), Larry J. Hillblom Foundation (2016-A-005-SUP [JSY]), Research Council of Norway (#213837, #225989, #223273, and #237250/EU JPND [OAA]), South East Norway Health Authority (2013-123), Norwegian Health Association, the Radiological Society of North America (RMS1741 [LWB]) (RSD), ASNR Foundation AD Imaging Award (RSD), National Alzheimer’s Coordinating Center (NACC) Junior Investigator (JI) Award (RSD), and the Tau Consortium (JSY).

